# Estrogens dynamically regulate neurogenesis in the dentate gyrus of adult female rats

**DOI:** 10.1101/2022.09.30.510371

**Authors:** Shunya Yagi, Yanhua Wen, Ariel A Batallán Burrowes, Samantha A Blankers, Liisa AM Galea

## Abstract

Estrone and estradiol differentially modulate neuroplasticity and cognition, but how they influence maturation pathways of new neurons in the adult hippocampus is not known. The present study assessed the effects of estrone and estradiol on various aspects of neurogenesis in the dentate gyrus (DG) of ovariectomized young adult Sprague-Dawley rats using daily subcutaneous injections of 17β-estradiol or estrone. Rats were injected with a DNA synthesis marker, 5-bromo-2-deoxyuridine (BrdU), and were perfused one, two, or three weeks after BrdU injection and treatment. Immunofluorescent labelling for GFAP/Sox2 was used to examine the density of neural stem/progenitor cells and Ki67 for cell proliferation. Double-immunofluorescent labelling of BrdU with doublecortin (DCX) or NeuN was used to examine the attrition and maturation of adult-born neurons over time. Length of time post ovariectomy was associated with a reduction in neural stem/progenitor cells in the DG. Estrogens had effects on different components of neurogenesis after one week of exposure but not after prolonged treatment. Estradiol enhanced, whereas estrone reduced, cell proliferation after one week but not after longer exposure to hormones. Both estrogens increased the density of BrdU/DCX-ir cells after one week of exposure but showed greater attrition of new neurons between one and two weeks after exposure, suggesting that the effects of estrogens on neurogenesis were not sustained. These results demonstrate that estrogens modulate several aspects of adult hippocampal neurogenesis differently in the short term, but may lose their ability to influence neurogenesis after long-term exposure.

## 1. Introduction

Hippocampal integrity is compromised in neurodegenerative and psychiatric disorders such as Alzheimer’s disease and depression (Scheff et al., 2006; Selden et al., 1991). A unique characteristic of the hippocampus is its ability to generate new neurons in adulthood. Neurogenesis in the adult hippocampus has important roles in pattern separation, during memory encoding, and in stress resilience (Anacker et al., 2018; Clelland et al., 2009; Tobin et al., 2019). New granule cells in the dentate gyrus (DG) of the hippocampus are produced from neural stem/progenitor cells in the subgranular zone. These cells share many morphological and functional similarities with radial glial cells and can develop into neurons, glia or remain undifferentiated. Neurons that develop from neural stem/progenitor cells express unique endogenous markers at different stages of development (Von Bohlen und Halbach, 2011; Kempermann et al., 2015). In the early stages, neural stem/progenitor cells express GFAP and Sox2, two proteins that are also individually found in mature astrocytes. However, when expressed together in stem/progenitor cells, GFAP/Sox2 are markers for putative neural stem/progenitor cells that will develop into new neurons (Boldrini et al., 2018). Ki67 is expressed in proliferating cells, immature neurons express doublecortin (DCX), and mature neurons express neuronal nuclei (NeuN), the timeline of which varies by sex and species (Snyder et al., 2009; Yagi et al., 2020). Although there are no sex differences in the number of new three-week-old neurons in rats, there are sex differences in the maturation pathways for adult neurogenesis (Yagi et al., 2020). Male rats have a greater density of neural stem/progenitor cells, greater cell proliferation, and faster maturation of new neurons than female rats (Yagi et al., 2020). However, male rats also have greater attrition of immature neurons between one and two weeks after production compared to female rats (Yagi et al., 2020). These findings suggest it may be fruitful to determine whether ovarian hormones, such as estrogens, regulate the maturation of neurogenesis in female rodents.

Estrone and estradiol are the two most abundant estrogens seen outside of pregnancy (Acconcia & Marino, 2018). Estradiol binds with greater affinity to estrogen receptors (ERs) (Kuiper et al., 1997) and is present at higher levels than estrone before menopause whereas estrone is present at higher levels than estradiol after menopause in human females (Rannevik et al., 1995). Previous studies demonstrate that these estrogens modulate neurogenesis in the DG depending on the type of estrogens, duration of exposure and component of neurogenesis (cell proliferation or survival of new neurons; Barha et al., 2009; Ormerod et al., 2003; McClure et al., 2013; Tanapat et al., 2005; Tanapat et al., 1999). Although both estrogens can increase cell proliferation, estradiol increases cell proliferation at multiple doses (Barha et al., 2009), estradiol enhances, whereas estrone decreases, the survival of three-week-old new neurons in rats that underwent cognitive training (McClure et al., 2013). In addition, acute exposure to estradiol enhances, whereas estrone impairs, contextual fear conditioning in adult female rats (Barha & Galea, 2010). Acute 17β-estradiol or estradiol benzoate (EB) rapidly increases cell proliferation (Barha et al., 2009; Ormerod et al., 2003; Mazzucco et al., 2006; Tanapat et al., 1999) but repeated administration of 17β-estradiol or EB for three weeks has no significant effect on cell proliferation (Tanapat et al., 2005; McClure et al., 2013; Chan et al., 2014). Collectively, these studies suggest that the different estrogens and duration of exposure can have distinct effects on various aspects of neurogenesis in the DG.

Estrogens also influence cognition and hippocampal volume in humans depending on the type, timing, and duration of hormone therapy (reviewed in Wnuk et al., 2012; Maki et al., 2009). Short duration of hormone therapy increases hippocampal volume (Erickson et al., 2007; Boyle et al., 2021), whereas longer than ten years of hormone therapy decreases hippocampal volume in postmenopausal females (Erickson et al., 2007). Furthermore, estradiol-based hormone therapy improves verbal memory in post-menopausal females, whereas estrone-based hormone therapy has detrimental (or non-significant) effects on verbal memory (Phillips & Sherwin, 1992; Shaywitz et al., 2003; Joffe et al., 2006; Linzmayer et al., 2001; Maki et al., 2007; Ryan et al., 2012). The timing of initiation of hormone therapy relative to menopause also plays an important role in the effects of hormone therapy, as early initiation after menopause enhances cognitive performance, whereas late initiation after menopause leads to poorer performance (MacLennan et al., 2006), an effect mirrored in animal models (reviewed in (Daniel & Bohacek, 2010).

To date, duration-dependent changes in the effects of estrogens have not been investigated on the characteristics of neurogenesis in the DG. Thus, we aimed to elucidate the effects of estrone and estradiol on neural stem/progenitor cells, the maturation rate of new neurons, and the trajectory (attrition) of new neurons. We hypothesized that estradiol and estrone would differentially modulate the trajectory and maturation rate of new neurons based on the duration of exposure to estrogens.

## 2. Materials and Methods

### 2.1. Subjects

Thirty-six female Sprague-Dawley rats (four females for each treatment and maturation time course) obtained from our breeding colony at the University of British Columbia (Vancouver, BC, Canada) were used in this study. Rats were weaned at postnatal day 21 and housed with same-sex siblings until puberty. Rats were then pair-housed until the end of the study in opaque polysulfone bins (432 mm × 264 mm × 324 mm) with paper towels, a single polycarbonate hut, virgin hardwood chip bedding, and free access to food and water. The colony room was maintained under a 12:12-h light/dark cycle (lights on at 07:00 h). All experiments were carried out in accordance with the Canadian Council for Animal Care guidelines and were approved by the animal care committee at the University of British Columbia. All efforts were made to reduce the number of animals used and their suffering during all procedures.

### 2.2. Experimental timeline

At 10-weeks-old, rats were handled for 2 minutes every day for two weeks. At 12-weeks-old, rats received a bilateral ovariectomy and were randomly assigned to one of three treatment groups. Following a one-week recovery period, rats received daily subcutaneous injections of 10 μg estrone (Sigma-Aldrich) in 0.1 ml sesame oil, 10 μg of 17β-estradiol (Sigma-Aldrich) in 0.1 ml sesame oil, or 0.1 ml sesame oil (vehicle). Daily subcutaneous injections of 10 μg of 17β-estradiol result in serum concentrations of estradiol that are equivalent to the proestrous phase but can also be supraphysiological (Woolley and McEwen, 1993). These doses of estrone and estradiol were chosen as they both result in increased levels of cell proliferation (Barha et al., 2009). Each group received the same treatment every day (approximately 9-11 am) until the end of the experiment. On the next day, all rats received one injection of bromodeoxyuridine (BrdU; 200 mg/kg i.p.; BD Biosciences) one hour after hormone or vehicle treatment. Rats were perfused one, two, or three weeks after BrdU injection (Figure 1A). Serum estradiol levels were 1.59 times higher in the estradiol group versus the estrone group, which was 50x and 20x higher than the oil-injected groups (verified via a multiplex electrochemiluminescence immunoassay kit (Custom Steroid Hormone Panel, Human/Mouse/Rat) from Meso Scale Discovery (Rockville, MD, USA)).

**Fig. 1.**
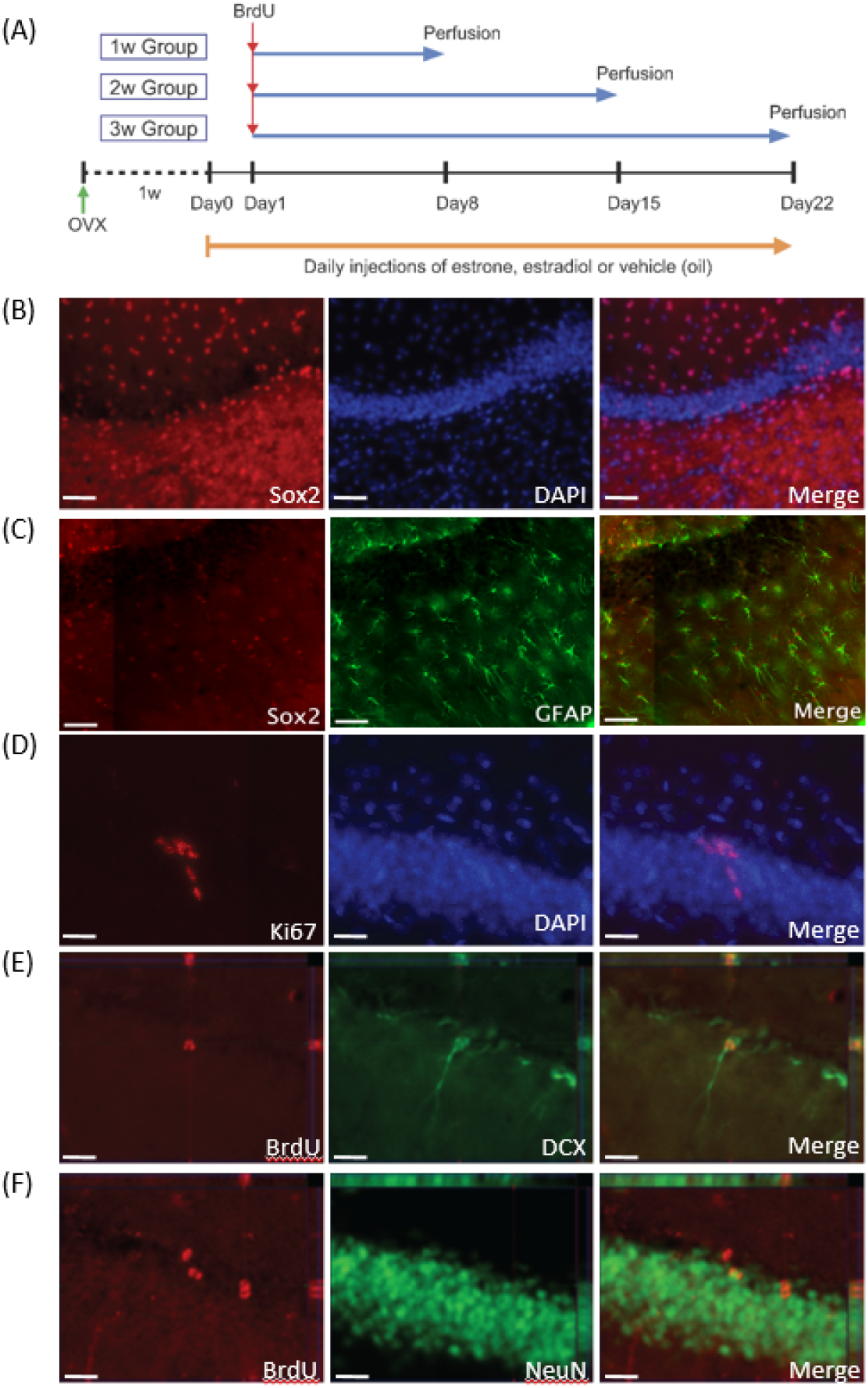
**A**: Schematic illustration for the experimental timeline. **B-E**: Photomicrographs for (B) Sox2-ir cells (red) with DAPI (blue), (C) Ki67-ir cells (red) with DAPI (blue), (D) BrdU-ir cells (red) with DCX-ir cells (green) and (E) BrdU-ir cells (red) with NeuN-ir cells (green). Scale bars in (B) indicate 50 μm and scale bars in (C)-(E) indicate 20 μm. All photomicrographs were taken by Zeiss Axio Scan.Z1 with 20x (B) or 40x (C-E) objectives.

### 2.3. Perfusion and tissue processing

Rats were administered an overdose of sodium pentobarbital (500 mg/kg, i.p.) and perfused transcardially with 60 ml of 0.9% saline followed by 120 ml of 4% formaldehyde (Sigma-Aldrich). Brains were extracted and post-fixed in 4% formaldehyde overnight, then transferred to 30% sucrose (Fisher Scientific) in solution for cryoprotection and remained in the solution until sectioning. Brains were sliced into 30 μm coronal sections using a Leica SM2000R microtome (Richmond Hill, Ontario, Canada). Sections were collected in series of ten throughout the entire rostral-caudal extent of the hippocampus and stored in an anti-freeze solution consisting of ethylene glycol (Sigma-Aldrich), glycerol (Avantor), and 0.1M PBS at –20°C.

### 2.4. Immunohistochemistry

Brain sections were stained for Sox2 (Figure 1B) or double-stained for Sox2/GFAP to examine the density of neural stem/progenitor cells, and Ki67 (Figure 1C) for the density of proliferating cells in the DG. Furthermore, brain sections were double-stained for BrdU/DCX (Figure 1D) and BrdU/NeuN (Figure 1E) to examine the maturation time course of new cells.

#### 2.4.1. BrdU/NeuN or BrdU/DCX double-labeling

Brain sections were prewashed three times with 0.1 M PBS and left overnight at 4°C. The tissue was incubated in a primary antibody solution containing 1:250 mouse anti-NeuN (Millipore Sigma; MA, USA) or 1:200 goat anti-DCX (Santa Cruz Biotechnology, CA, USA), 0.3% Triton-X (Sigma-Aldrich), and 3% normal donkey serum (NDS; Vector Laboratories) in 0.1 M PBS for 24 hours at 4°C. Following three rinses in 0.1 M PBS, sections were incubated in a secondary antibody solution containing 1:200 donkey anti-mouse Alexa Fluor 488 (Invitrogen, Burlington, ON, Canada) or 1:200 donkey anti-goat Alexa Fluor 488 (Invitrogen, Burlington, ON, Canada) in 0.1 M PBS, for 18 hours at 4°C. After rinsing three times with PBS, the sections were washed with 4% formaldehyde, and rinsed twice in 0.9% NaCl (Sigma-Aldrich), followed by incubation in 2N HCl (Fisher Scientific) for 30 minutes at 37°C. Following three rinses in 0.1 M PBS, the sections were then incubated in a BrdU primary antibody solution consisting of 1:1000 rat anti-BrdU (AbD Serotec; Raleigh, NC, USA), 3% NDS, and 0.3% Triton-X in 0.1 M PBS for 24 hours at 4°C. Sections were then incubated in a secondary antibody solution containing 1:500 donkey anti-rat Cy3 (Jackson ImmunoResearch; PA, USA) in 0.1 M PBS for 24 hours at 4°C. Following three rinses with PBS, the sections were mounted onto microscope slides and cover-slipped with PVA DABCO.

#### 2.4.2. Sox2 and Sox2/GFAP double-labeling

Brain sections were prewashed with 0.1 M PBS and left to sit overnight at 4°C. The next day, sections were washed in 0.1M PBS for 10 min each and blocked with 3% NDS and 0.3% Triton X-100 in 0.1 M PBS, followed by incubation in primary antibody solution made with 1:1000 mouse anti-Sox2 (Santa Cruz Biotechnology), 1% NDS, and 0.3% Triton X-100 in 0.1 M PBS for 24 h at 4°C. Then the sections were incubated in a secondary antibody solution, consisting of 1:500 donkey anti-mouse Alexa Fluor 594 (Invitrogen), 1% NDS, and 0.3% Triton X-100 in 0.1 M PBS, for 18 h at 4°C. After three rinses with PBS, the sections were incubated in 1:5000 DAPI (Life Technologies) in PBS for 3 min. Followed by three rinses, tissues were mounted onto slides and cover-slipped with PVA DABCO.

For the double-labelling, brain sections were prewashed with 0.1 M PBS and left to sit overnight at 4°C. The next day, sections were washed in 0.1M PBS for 10 min each and blocked with 3% NDS and 0.3% Triton X-100 in 0.1 M PBS, followed by incubation in primary antibody solution made with 1:1000 mouse anti-Sox2, 1:500 rabbit anti-GFAP (Invitrogen), 1% NDS, and 0.3% Triton X-100 in 0.1 M PBS for 24 h at 4°C. Then the sections were incubated in a secondary antibody solution, consisting of 1:500 donkey anti-mouse Alexa Fluor 594, 1:500 donkey anti-rabbit Alexa Fluor 488 (Thermo Fisher Scientific), 1% NDS, and 0.3% Triton X-100 in 0.1 M PBS, for 18 h at 4°C. After three rinses with PBS, the sections were incubated in 1:5000 DAPI in PBS for 3 min. Followed by three rinses, tissues were mounted onto slides and cover-slipped with PVA DABCO.

#### 2.4.3. Ki-67

Brain sections were prewashed with 0.1 M PBS and left to sit overnight at 4°C. The next day, sections were incubated in 10 mM sodium citrate buffer (Sigma Aldrich) for 30 min at 90°C to retrieve antigens of Ki67 and blocked with 3% NDS and 0.3% Triton X-100 in 0.1 M PBS. Tissue was then incubated in primary antibody solution made with 1:250 mouse anti-Ki67 (Leica Biosystems), 1% NDS, and 0.3% Triton X-100 in 0.1 M PBS for 24 h at 4°C. Following three washes in 0.1 M PBS, brain sections were incubated in a secondary antibody solution, consisting of 1:500 donkey anti-mouse Alexa Fluor 488 (Invitrogen), 1% NDS, and 0.3% Triton X-100 in 0.1 M PBS, for 18 h at 4°C. After three rinses with PBS, sections were incubated in 1:5000 DAPI in PBS for 3 min. Followed by three rinses, tissue was mounted onto slides and cover-slipped with PVA DABCO.

### 2.5. Cell counting

All counting was conducted by an experimenter blind to the group assignment of each animal using an Olympus FV1000 confocal microscope and/or Zeiss Axio Scan.Z1 (Carl Zeiss Microscopy, Thornwood, NY, USA). The density of immunoreactive (ir) cells was calculated by dividing the total number of –ir cells by volume (mm^3^) of the corresponding region. Volume estimates were calculated by multiplying the summed areas by thickness of sections (0.03 mm, using Cavalieri’s principle; (Gundersen & Jensen, 1987). Area measurements for the region of interest were obtained using digitized images on Zen 3.0 software (blue edition; Carl Zeiss Microscopy, Thornwood, NY, USA). Cells were categorized as to whether they were in the dorsal or ventral DG using the criterion defined by Banasr and others (2006), with sections 6.20-3.70 mm from the interaural line defined as dorsal and sections 3.70-2.28 mm from the interaural line as ventral. Cells were counted separately in each region because the dorsal hippocampus is associated with spatial learning and memory, whereas the ventral hippocampus is associated more with stress and anxiety (Moser et al., 1993; Kjelstrup et al., 2002).

BrdU-ir cells were counted under a 60x oil immersion objective lens using an Olympus epifluorescent microscope. The percentage of BrdU/NeuN-ir cells were obtained by randomly selecting 50 BrdU-ir cells and calculating the percentage of cells that were double-labelled-ir with NeuN under 40x objective lens using an Olympus FV1000 confocal microscope (Olympus, Richmond Hill, ON, Canada). The percentages of BrdU/DCX-ir cells were obtained by randomly selecting 50 BrdU-ir cells and calculating the percentage of cells that double-labelled-ir with DCX on digitized images acquired under 40x objective lens using Axio Scan.Z1 slide scanner with Zen 3.0 software (blue edition; Carl Zeiss Microscopy, Thornwood, NY, USA).

Ki67-ir cells were counted on digitized images every twentieth section. Photomicrographs for Ki67-ir cells were taken with a 40x objective lens on an Axio Scan.Z1 slide scanner with Zen 3.0 software (Carl Zeiss Microscopy, Thornwood, NY, USA). Photomicrographs for Sox2-ir cells and Sox2/GFAP-ir were taken from four dorsal and three ventral hippocampal sections using a 40x objective lens on an Axio Scan.Z1 slide scanner, and optical density of Sox2-ir cells were measured on digitized images using ImageJ (NIH, Bethesda, MD, USA). The proportion of Sox2/GFAP-ir cells was obtained by randomly selecting 50 GFAP-ir cells and counting how many of them were double-labeled with Sox-2 on digitized images taken with an Axio Scan.Z1 slide scanner and 40x objective.

### 2.6. Statistical analyses

All analyses were conducted using STATISTICA (Statsoft Tulsa, OK). Repeated-measures analysis of variance (ANOVAs) were used to analyze the density of Ki67-ir, Sox2-ir, and BrdU-ir cells, and the percentage of Sox2/GFAP-ir. BrdU/DCX-ir, and BrdU/NeuN-ir cells with exposure time (1w, 2w, 3w) and hormone (estrone, estradiol, vehicle) as between-subject factors and hippocampal region (dorsal, ventral) as the within-subject factor. Post-hoc tests utilized the Neuman-Keuls procedure. A priori comparisons were subjected to Bonferroni corrections. Significance was set to α=0.05 and effect sizes are given with Cohen’s d or partial η^2^.

## 3. Results

### 3.1. Estradiol reduced the density of Sox2-ir cells in the dorsal DG, whereas estrone reduced the density of Sox2-ir cells in the ventral DG compared to vehicle-treated females

Both estrogens reduced the density of Sox2-ir cells dependent on region but not on exposure time. Estradiol-treated females had a lower density of Sox2-ir cells in the dorsal DG compared to vehicle-treated females (p = 0.021, Cohen’s d = 1.222), whereas estrone-treated females had a lower density of Sox2-ir cells in the ventral DG compared to both groups (vehicle-treated: p = 0.002, Cohen’s d = 1.410; estradiol-treated: p = 0.002, Cohen’s d = 1.000) [interaction effect of region by hormone: F(2, 25) = 5.60, p < 0.01, partial η^2^ = 0.309: Figure 2A and 2B]. Furthermore, there was a greater density of Sox2-ir cells in the ventral DG compared to the dorsal DG in estradiol-treated (p < 0.001, Cohen’s d = 1.909) and vehicle-treated females (p = 0.01, Cohen’s d = 0.959) but not in estrone-treated females (p=0.53). There was also a main effect of hormone [F(1, 25) = 4.19, p = 0.027, partial η^2^ = 0.251] and region as expected [F(1, 25) = 29.36, p < 0.001, partial η^2^ = 0.540]. There were no other significant main or interaction effects on the density of Sox2-ir cells (p > 0.180).

**Fig. 2.**
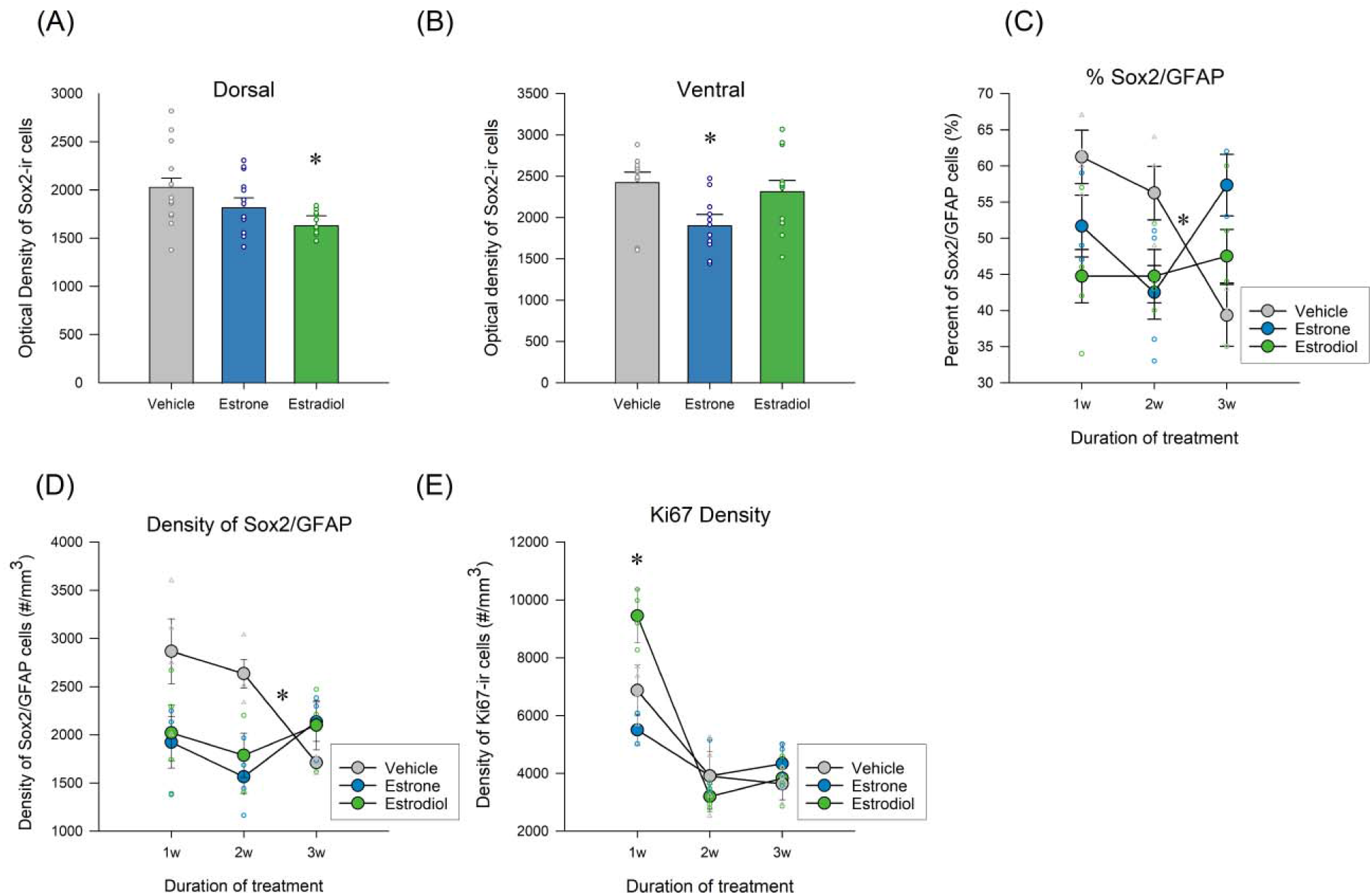
**A-B**: Mean (±SEM) density of Sox2-ir cells in the dorsal dentate gyrus (A) and the ventral dentate gyrus (B). Vehicle-treated females had a greater density of Sox2-ir cells compared to estradiol-treated females in the dorsal dentate gyrus, and estradiol-treated and vehicle-treated females had a greater density of Sox2-ir cells in the ventral dentate gyrus. **C-D**: Mean (±SEM) density of Ki67-ir cells in the dorsal dentate gyrus (C) and the ventral dentate gyrus (D). Estradiol-treated females showed greater density of Ki67-ir cells compared to vehicle-treated females and compared to estrone-treated females, and vehicle-treated females showed greater density of Ki67-ir cells compared to estrone-treated females after one week of hormone exposure. * indicates p < 0.05. SEM-standard error of the mean.

### 3.3. Ovariectomy decreased the proportion of neural stem/progenitors cells in the DG with time, whereas estrogens maintained stable proportions of neural stem/progenitor cells in the DG

The proportion of Sox2/GFAP-ir cells decreased in the vehicle treated group but was stable in the groups treated with estrogens. After three weeks of treatment, the proportion of Sox2/GFAP-ir cells in oil-treated females decreased in comparison to weeks one (p = 0.033, Cohen’s d = 2.325), but there was no significant change across weeks with estradiol (p’s >0.60) or estrone (p’s >0.62) treatment [interaction effect of hormone by week: F(4,24) = 5.165, p = 0.004, partial η^2^ = 0.463; Figure 2C]. There was also a main effect of region as expected [F(1,24) = 36.241, p < .001, partial η2 = 0.061]. There were no other significant main effects or interactions in the proportion of Sox2/GFAP-ir cells (all p’s > 0.157). When examining the density of Sox2/GFAP co-expressing cells, we found that both estrogens had reduced density of progenitor cells (main effect of hormone (F(4,23) = 4.356, p = 0.024, partial η^2^ = 0.274) and that only vehicle-treated females showed a reduction over time (oil p=0.002, Cohen;s d=4.52, but not the estrogens all p>0.5).

### 3.2. Estradiol increased, whereas estrone reduced, the density of Ki67-ir cells in the DG compared to vehicle-treated females after one week of hormone exposure

After one week of hormone treatment, estradiol-treated females had a greater density of Ki67-ir cells compared to both groups (vehicle-treated: p < 0.001, Cohen’s d = 2.864; estrone-treated: p < 0.001, Cohen’s d = 5.239), whereas estrone-treated females had a lower density of Ki67-ir cells compared to both groups (vehicle-treated: p = 0.026, Cohen’s d = 1.883) [interaction effect of exposure time by treatment: F(4, 26) = 10.453, p < 0.001, partial η^2^ = 0.617: Figure 2E]. Furthermore, the density of Ki67-ir cells was highest at one, regardless of group [all p’s < 0.05]. There were also main effects of hormone [F(2, 26) = 4.061, p = 0.029, partial η^2^ = 0.238], week [F(2, 26) = 70.034, p < 0.001, partial η^2^ = 0.843] and region [F(1, 26) = 18.033, p < 0.001, partial η^2^ = 0.410]. There were no other significant main or interaction effects on the density of Ki67-ir cells (all p’s > 0.608).

### 3.3. Estrone– and estradiol-treated females had a greater density of BrdU-ir cells and BrdU/DCX-ir cells compared to vehicle-treated females one week after exposure to estrogens

After one week of exposure to estrogens, estrone-treated (p = 0.002, Cohen’s d = 2.655) and estradiol-treated females (p = 0.004, Cohen’s d = 1.868) had a greater density of one-week-old BrdU-ir cells compared to vehicle-treated females [interaction effect of week by hormone: F(4, 24) = 3.120, p = 0.034, partial η^2^ = 0.342: Figure 3A], but there were no significant differences between treatment groups after two or three weeks of hormone treatment in BrdU-ir cells (p’s > 0.184). In both estrone and estradiol-treated females, there was a significantly greater density of BrdU-ir cells at one week compared to two weeks (estrone: p = 0.004, Cohen’s d = 3.955; estradiol: p = 0.001, Cohen’s d = 2.980) but no significant difference between two weeks and three weeks (p’s > 0.118). In vehicle-treated females, there were no significant differences in the density of BrdU-ir cells between any of the weeks (p’s > 0.302). There were also significant main effects of hormone [F(2, 24) = 7.550, p = 0.003, partial η^2^ = 0.342], week [F(2, 24) = 30.114, p < 0.001, partial η^2^ = 0.715] and region [F(1, 24) = 4.284, p = 0.049, partial η^2^ = 0.151], but no other significant interaction effects on the density of BrdU-ir cells (p’s > 0.288).

**Fig. 3.**
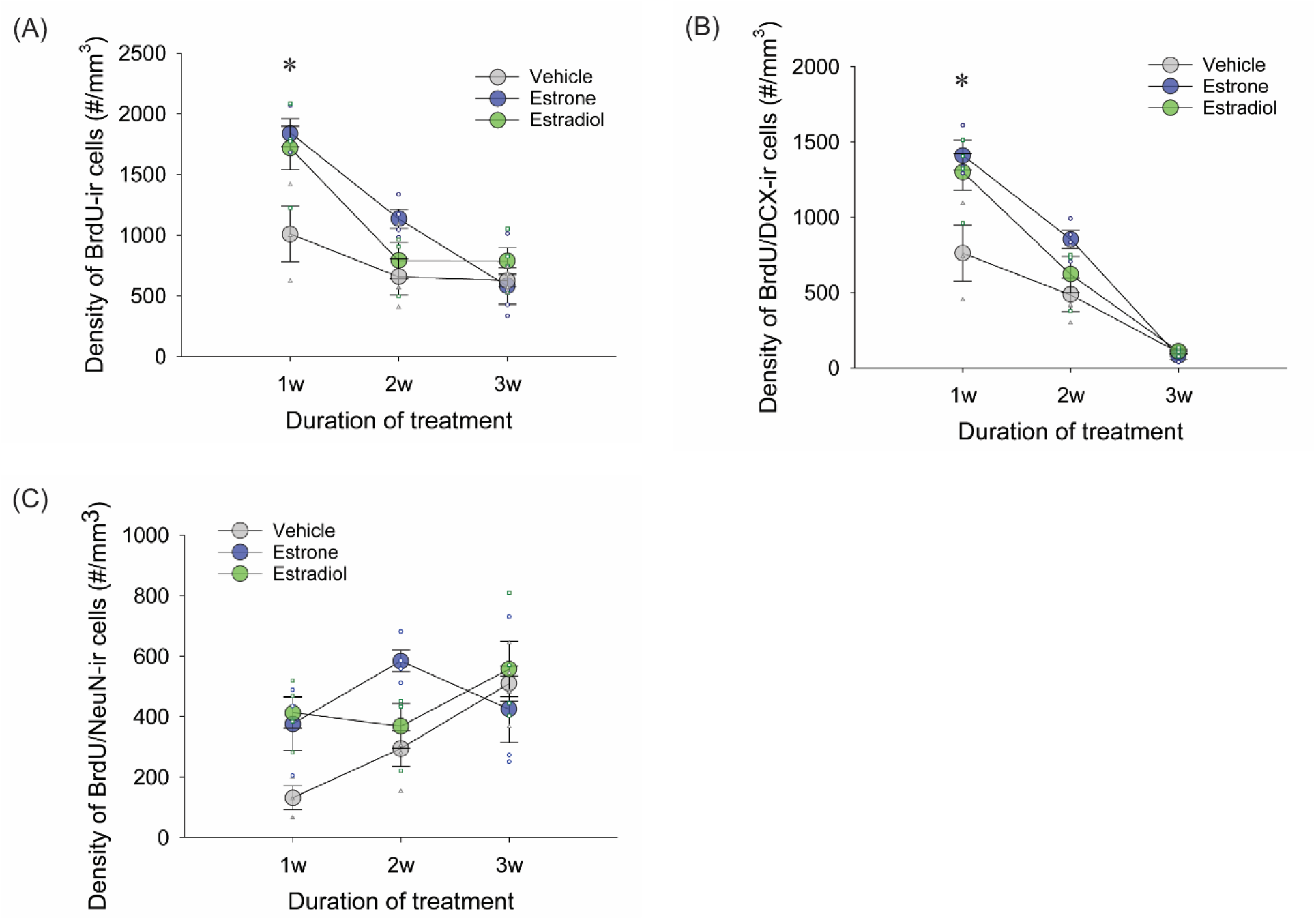
**A**: Mean (±SEM) density of BrdU-ir cells in the dentate gyrus. **B:** Mean (±SEM) density of BrdU/DCX-ir cells in the dentate gyrus. Estrone or estradiol-treated females had a greater density of BrdU-ir cells and BrdU/DCX-ir cells in the ventral dentate gyrus compared to vehicle-treated females one week after BrdU injection and exposure to hormones. **C:** Mean (±SEM) density of BrdU/NeuN-ir cells in the dentate gyrus. Estradiol-treated females had a trend of greater density of BrdU/NeuN-ir cells compared to vehicle-treated females one week after BrdU injection and exposure to hormones. D: Mean (± SEM) percentage of Sox2/GFAP-ir cells in the dorsal dentate gyrus. Estrogen-treated females had stable proportions of Sox2/GFAP-ir cells across the three weeks, compared to oil-treated females that had stable proportions of cells in weeks one and two, but decreased in week three. * indicates p < 0.05. SEM-standard error of the mean.

One week of exposure to estrogens also increased the density of BrdU/DCX-ir cells (estrone: p < 0.001, Cohen’s d = 2.529; estradiol: p = 0.001, Cohen’s d = 1.903) compared to vehicle-treated females [interaction effect of hormone by week: F(4, 24) = 4.109, p = 0.011, partial η^2^ = 0.406: Figure 3B]. As expected the density of BrdU/DCX-ir cells was decreased with across weeks [one to two weeks: p’s < 0.001, Cohen’s d = 3.777 (estrone), Cohen’s d = 3.026 (estradiol); two to three weeks: p’s < 0.002, Cohen’s d = 8.906 (estrone), Cohen’s d = 3.430 (estradiol); vehicle-treated females (one to two weeks: p = 0.103; two to three weeks: p = 0.019, Cohen’s d = 2.403). There was also a significant main effect of hormone [F(2, 24) = 10.249, p < 0.001, partial η^2^ = 0.461] and week [F(2, 24) = 101.756, p < 0.001, partial η^2^ = 0.895]. There were no other significant main or interaction effects on the density of BrdU/DCX-ir cells (p > 0.167).

In terms of the density of mature new neurons over the weeks, only the vehicle-treated groups that showed a significant increase in the density of BrdU/NeuN-ir cells across time from one to three weeks (p=0.018, Cohen’s d = 3.974) which was not seen in the groups treated with estrogens (p > 0.401; interaction effect of hormone by week: F(4, 24) = 2.963, p = 0.040, partial η^2^ = 0.331; Figure 3C). This was likely due to earlier maturation of new neurons by estrogens as estradiol-treated and estrone-treated females had a trend for a greater density of BrdU/NeuN-ir cells compared to vehicle-treated females after one week (estradiol: p = 0.077, Cohen’s d = 3.225; estrone: p = 0.095). There were also main effects of hormone [F(2, 24) = 3.932, p = 0.033, partial η^2^ = 0.247] and week [F(2, 24) = 4.963, p = 0.016, partial η^2^ = 0.293]. There were no other main or interaction effects on the density of BrdU/NeuN-ir cells (all p’s > 0.208).

### 3.4. Shorter durations of estradiol exposure increased percentage of BrdU cells that also co-expressed immature or mature neuronal proteins

Despite a significant interaction effect of hormone by week, there were no differences among groups within each week, except that estradiol-treated females tended to have a greater percentage of BrdU/DCX-ir cells in the DG compared to vehicle-treated females at the two week time point [p =0.067, Cohen’s d = 4.139; interaction effect of hormone by week: F(4, 25) = 2.96, p = 0.040; Figure 4A and 4B], but not after one week or three weeks of hormone treatment/after BrdU injection (p’s > 0.188). Furthermore, the percentage of BrdU/DCX-ir cells was greater in the dorsal DG compared to the ventral DG after two weeks [interaction effect of week by region: F(2, 25) = 5.04, p = 0.015, partial η^2^ = 0.287; post-hoc: p = 0.011, Cohen’s d = 1.249]. There were main effects of week [F(2, 25) = 2467.46, p < 0.001, partial η^2^ = 0.995] and region [F(1, 25) = 4.62, p = 0.042, partial η^2^ = 0.156]. There were no other significant main or interaction effects on the percentage of BrdU/DCX-ir cells (p > 0.306).

**Fig. 4.**
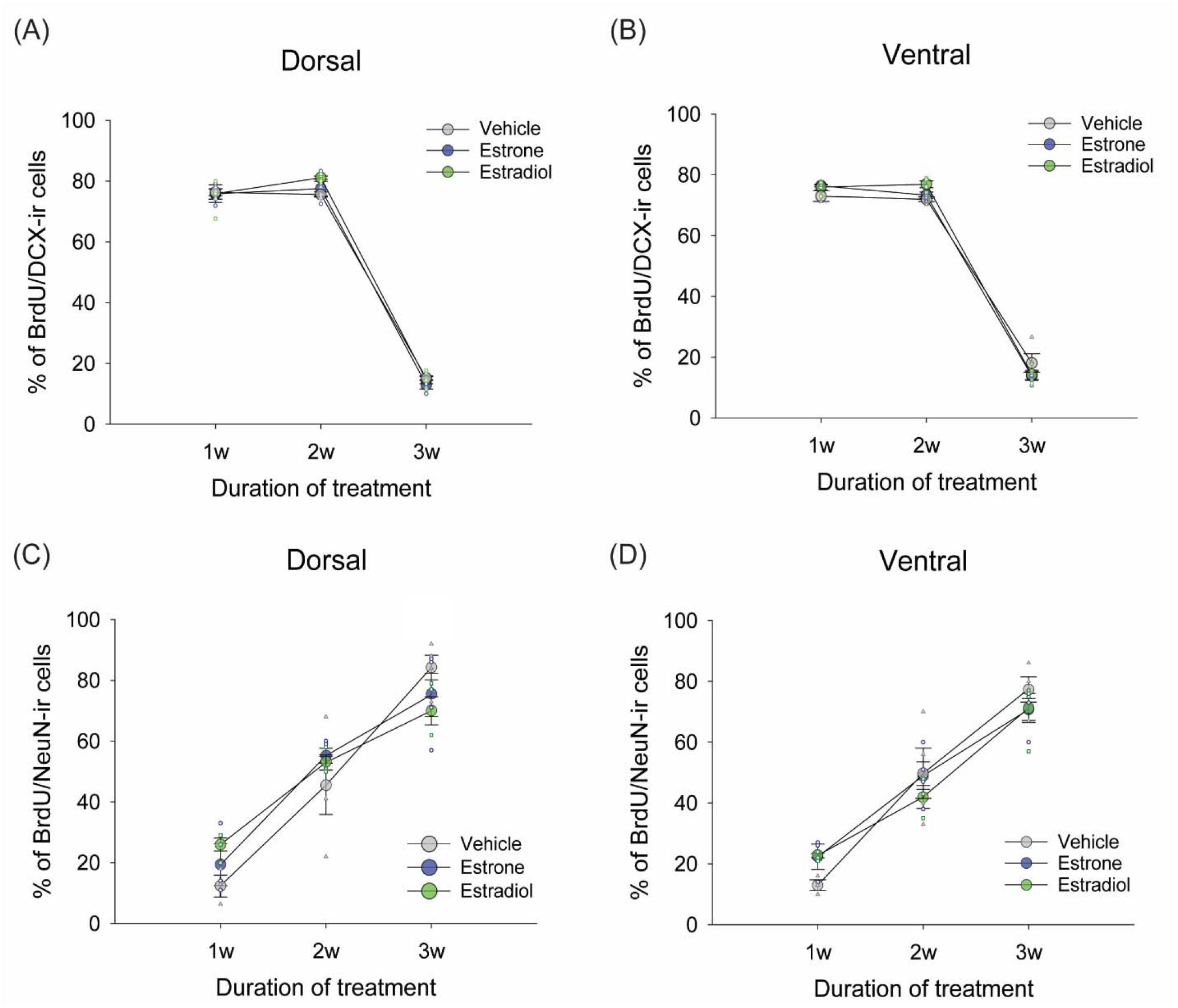
**A-B**: Mean (±SEM) percentage of BrdU/DCX-ir cells in the dorsal dentate gyrus (A) and the ventral dentate gyrus (B). Estradiol-treated female rats showed a trend of greater percentage of BrdU/DCX-ir cells compared to vehicle-treated female rats two weeks after BrdU injection. **C-D**: Mean (±SEM) percentage of BrdU/NeuN-ir cells in the dorsal dentae gyrus (C) and the ventral dentate gyrus (D). Estradiol-treated females, compared to vehicle-treated females, showed a greater percentage of BrdU/NeuN-ir cells in the dorsal DG three weeks after BrdU injection/hormone exposure. * indicates p < 0.05. SEM-standard error of the mean.

As expected, the percentage of BrdU/NeuN-ir neurons increased across weeks [main effect of week: F(2, 24) = 104.616, p < 0.001, partial η^2^ = 0.897: Figure 4C-D]. Based on the findings above on the density of BrdU/NeuN or BrdU/DCX cells we expected that one week of estradiol would increase the percentage of BrdU that were co-labelled with NeuN, and this comparison was significant (one-tailed p= 0.049, one-tailed, Cohen’s 2.43). There was a trend for a significant main effect of region [F(1, 24) = 3.136, p = 0.089, partial η^2^ = 0.116] and an interaction of region by hormone by week [F(2, 24) = 2.230, p =0.096, partial η^2^ = 0.271], but no other significant main or interaction effects on the percentage of BrdU/NeuN-ir cells in the DG (p > 0.096).

## 4. Discussion

Both estrogens had a dynamic effect on different characteristics of neurogenesis which depended on the duration of exposure to estrogens. Furthermore, longer time post ovariectomy reduced the density of neural stem/progenitor cells in the DG over time, whereas exposure to estrogens eliminated this affect, suggesting a potential mechanism for why estrogens may lose their potency to alter neuroplasticity with longer withdrawal periods post ovariectomy. Estradiol increased cell proliferation after a week of exposure but estrone decreased cell proliferation after one week of exposure . Shorter exposure to estrogens (one week) increased the density of new neurons and enhanced the early maturation of new neurons compared to vehicle exposure. However, a longer duration of exposure to estrogens (2-3 weeks) resulted in greater attrition of immature neurons such that there was no longer a significant difference in the number of new neurons after three weeks of exposure to estrogens compared to controls. These findings suggest that the pathways to alter neurogenesis differ by the type of estrogens and duration of exposure, which reflects early but not sustained neurogenic properties of estrogens. These findings highlight the importance of studying estrogen type and duration in females, which have important implications for treatments that promote hippocampal plasticity.

### 4.1. As the time since ovariectomy increased, there was a reduction in the expression of neural stem/progenitor cells in the DG

In the present study, vehicle-treated females showed a decrease in the proportion of Sox2/GFAP-ir cells three weeks after OVX compared to the first week. These findings indicate that prolonged deprivation of ovarian hormones results in a decline in neural progenitor cells by 4 weeks post ovariectomy (approx. 20%). Comparatively, estrogens maintain stable levels of neural stem/progenitor cells across time although at least with the density of Sox2/GFAP, this was due to a decrease with estrogens in the density within the first week. This result is intriguing as it suggests that the longer time post-ovariectomy the fewer stem cells would be able to be stimulated into new neurons. This may be one mechanism by which hormone therapy is no longer effective with more time post menopause. Recent meta-analyses suggest that late-life HT is associated with increased risk of dementia compared to closer to menopause (Nerattini et al., 2023). Animal models also suggest that estrogens are not as effective after long–term ovariectomy compared to shortly after ovariectomy ( Bohacek et al., 2008; Ma et al., 2020; Wu et al., 2011).

One week of exposure to estradiol and estrone reduced the density of neural stem/progenitor cells in the DG compared to vehicle, indicated by reduced levels of Sox2-ir. Interestingly, the effect of estrogens on neural stem/progenitor cells varied along the dorsoventral axis, where estradiol reduced neural stem/progenitor cells in the dorsal region and estrone in the ventral region. The dorsoventral axis of the hippocampus has differing functions as the dorsal region plays an important role in reference memory, and the ventral region plays an important role in stress, anxiety, and working memory (reviewed in Leary & Cryan, 2014). Previously, we found that intact females had a greater density of Sox2-ir cells in the ventral DG compared to dorsal DG (Yagi et al., 2020) and this same dorsoventral axis difference was seen in vehicle-treated and estradiol-treated females, but not in estrone-treated females. Underlying mechanisms or functional consequences for the differential regulation of estrogens on neural stem/progenitor cells along the dorsoventral axis have yet to be determined.

### 4.2. After one week of hormone treatment, estradiol enhances, whereas estrone reduces, cell proliferation in the DG

Estradiol enhanced cell proliferation in the DG after one week of treatment, but not after two or three weeks of hormone exposure. Interestingly, estrone had the opposite effect as it reduced cell proliferation after one week of exposure. These results suggest that estradiol, but not estrone, enhances cell proliferation in a limited time window (up to one week) in naïve rats. These results are consistent with previous work demonstrating that a single dose of estradiol enhances cell proliferation in the DG, whereas three weeks of repeated administration of estradiol had no significant effects on cell proliferation (Barha et al., 2009; Ormerod et al., 2003; Tanapat et al., 2005; McClure et al., 2013; Chan et al., 2014). In contrast, the present study found that estrone decreased cell proliferation in the DG after one week of treatment indicating that chronic estrone treatment has detrimental effects on cell proliferation in the short term. Previous work showed that the single injection of the same dose of estrone rapidly enhanced cell proliferation (Barha et al., 2009), whereas three weeks of repeated administration of estrone had no significant effects on cell proliferation (McClure et al., 2013). These findings indicate that estrone has dynamic effects on cell proliferation with an initial increase followed by a reduction and finally no influence on cell proliferation, possibly via a compensatory effect. The same dose of estradiol rapidly increases cell proliferation (at 30min), then reduces cell proliferation (48 h) but at 1 week increases cell proliferation but ultimately not affecting cp after three weeks of treatment. Why these different estrogens alter cell proliferation in distinct ways is not known. Ultimately for both estrogens after three weeks, there is no longer any significant influence on cell proliferation, suggesting compensatory actions of the progenitor cells, and future studies should be conducted to determine why these estrogens are no longer stimulatory after prolonged treatment.

### 4.3. Estrone and estradiol increased the density of one-week-old neurons after one week of exposure and increased the attrition of new neurons between one and two weeks

Despite the differential effects of estrone versus estradiol on cell proliferation after one week of exposure, both estrogens increased the density of one-week-old new cells (BrdU-ir cells) compared to vehicle treatment, suggesting that there are different modulatory pathways of estradiol and estrone on neurogenesis. However, the effect of prolonged estrogens on the survival of new BrdU-ir cells was not maintained as there was significant attrition of these new cells after three weeks of hormone treatment. As the majority of BrdU-ir cells (80%) at one and two weeks expressed an immature neuronal marker (DCX), the attrition of BrdU-ir cells was most likely due to the reduction of immature neurons. It is possible that evaluating a time point earlier than one week would have revealed a greater attrition of BrdU/DCX-ir cells as we saw the largest density of BrdU/DCX-ir cells 24 hours after BrdU injection in intact females (Yagi et al., 2020). Indeed, the percentage of BrdU-ir cells expressing NeuN was greater in the E2-treted group at one week, suggesting faster maturation of these new cells into neurons, though this maturation advantage did not translate into a greater density of new neurons at three weeks, again pointing to the greater attrition. It is possible with stimulation, these new neurons could be “rescued” from attrition in the “use it or lose it” fashion. Indeed, in rats undergoing Morris Water maze training during 3 weeks of treatment of estradiol at the same dose, there was an increased level of neurogenesis with E2 but a decrease with E1. Therefore, further research examining the functional characteristics of these immature neurons is required to make a solid conclusion on the effects of estrogens on the maturation of new neurons. Thus in naïve rats, these results indicate that although estrone and estradiol initially enhance adult neurogenesis (BrdU/DCX-ir and a trend for BrdU/NeuN-ir) at one week compared to vehicle treatment, there is no pro-neurogenic effect of estrogens in the long-term due to significant attrition in these new neurons with exposure to estrogens across time.

### 4.4. Implications

Our results suggest that both the type of estrogens and duration of exposure to estrogens can significantly influence neurogenesis in the hippocampus. These findings are interesting as both animal and human studies suggest duration of exposure to estrogens influences a variety of factors. Estrogen exposure has differential effects on cell proliferation depending on time since ovariectomy surgery as estrogens enhance cell proliferation after short-term (one week) ovarian hormone depletion (Barha et al., 2009; Ormerod et al., 2003; Mazzucco et al., 2006; Tanapat et al., 2005; Tanapat et al., 1999), whereas estrogens do not significantly influence cell proliferation after long-term (four weeks) depletion (Tanapat et al., 2005). Here we found that one week of estradiol exposure, but not longer-term exposure, enhanced both cell proliferation and maturation of new neurons after one week. Our findings are reminiscent of findings from hormone therapy studies, as early initiation of hormone therapy relative to menopause increases hippocampal volume, whereas late treatment initiation has no such beneficial effects on hippocampal volume (Erickson et al., 2010). In addition, we and others have found that the type of estrogens used can have opposing effects on neuroplasticity in human and animal studies(McClure et al., 2013; Phillips & Sherwin, 1992; Shaywitz et al., 2003; Joffe et al., 2006; Linzmayer et al., 2001; Maki et al., 2007; Boccardi et al., 2006; Resnick et al., 2009). Here, we found that one week of exposure to estradiol enhanced cell proliferation but one week of exposure to estrone decreased cell proliferation. This is consistent with findings that estradiol enhanced the survival of new neurons after three weeks of exposure, whereas estrone decreased neuron survival, in rodents that underwent cognitive training in the Morris water maze (McClure et al., 2013). Indeed, estradiol-based hormone therapy improves verbal memory and increases hippocampal volume (Phillips & Sherwin, 1992; Joffe et al., 2006; Boccardi et al., 2006), whereas conjugated estrone-based hormone therapy can have a detrimental effect in post-menopausal women(Phillips & Sherwin, 1992; Maki et al., 2007; Resnick et al., 2009). Thus, in humans and in rodents, different estrogens modulate neuroplasticity and cognition depending not only on the initiation of treatment relative to menopause/ovariectomy, but also on the type of hormone therapy (Barha et al., 2009; Ormerod et al., 2003; Tanapat et al., 2005; Tanapat et al., 1999; McClure et al., 2013; Chan et al., 2014; Phillips & Sherwin, 1992; Shaywitz et al., 2003; Maki et al., 2007; Boccardi et al., 2006; Resnick et al., 2009; Barker & Galea, 2008). Future studies need to take into account not only the type of hormone therapy but the time of menopause and relation of treatment to menopause.

## 5. Conclusion

Here we report that estrogens influence different facets of neurogenesis depending on the type of estrogen and duration of exposure. Our findings add to the growing literature that estrone and estradiol have similar but not equivalent effects on neurogenesis. We also show that the duration of exposure to estrogens has dynamic effects on neurogenic parameters, with proneurogenic effects within one week of exposure that are no longer evident with prolonged exposure to estrogens. Our findings shed light on the importance of studying the short– and long-term consequences of exogenous estrogens on adult neurogenesis.

## Acknowledgments

We would like to thank Jared Splinter, Stephanie Lieblich and Tallinn Splinter for the exceptional technical assistance with this work.

## Statement of Ethics

All experiments were carried out in accordance with the Canadian Council for Animal Care guidelines and were approved by the animal care committee at the University of British Columbia. All efforts were made to reduce the number of animals used and their suffering during all procedures.

## Author Contributions

**SY:** Conceptualization, Methodology, Data Collection, Data curation, Writing-Original draft preparation, Visualization. **YW**: Data collection. **ABB**: Data analysis, Draft revision. **LG**: Conceptualization, Methodology, Analysis, Writing-Reviewing and Editing, Supervision.

## Data Availability

The datasets used and/or analyzed during the current study are available from the corresponding author on reasonable request.

